# Landscape structure and climate seasonality affect the amount, richness and diversity of pollen collected by honeybees in a Neotropical region of Colombia

**DOI:** 10.1101/2020.05.01.072082

**Authors:** Paula María Montoya-Pfeiffer, Guiomar Nates-Parra

**Author notes:** **Corresponding author:** (PMMP).

## Abstract

Pollen is the main food for honeybee broods and young workers and so colony development and reproduction rely heavily on pollen availability, both spatially and temporally, in the environment. Intensification of agriculture and climate seasonality are known to alter honeybee foraging patterns and pollen intake through changes in resource availability in temperate regions; however, little is known about how honeybees respond to such environmental factors in tropical regions.

Pollen species collected by honeybees in a Neotropical agricultural region of Colombia were identified. The effects of landscape structure (landscape Shannon Diversity Index, forest area in 1000 m around the apiary) and climate seasonality (mean monthly precipitation) on the amount, richness and diversity of pollen collected by the honeybees were evaluated for all pollen species together and pollen species segregated according to forest and anthropic areas (croplands, grasslands, woodlands, urban areas).

Honeybees were found to be much more associated with anthropic than forest pollen species regardless of landscape structure or precipitation. However, the amount, richness and diversity of pollen from all species and forest species responded positively to landscape diversity and forest area, suggesting an advantage for honeybees in obtaining small quantities of pollen from forest species, in spite of being well-adapted to forage in anthropic areas. Precipitation was found not to be related to the overall amount and overall richness of pollen collected by honeybees, suggesting that climate seasonality was not an important factor for pollen foraging. Nonetheless, overall pollen diversity was negatively affected by precipitation in less diverse landscapes, while anthropic pollen diversity was negatively affected in more forested landscapes. These findings are compared with those from temperate regions, and the implications for honeybee productivity and survival, and their interactions with Neotropical native species, are discussed.

## Introduction

Pollen is essential for the development and sustainment of honeybee colonies because it provides the proteins necessary for the development of broods and young workers [1]. The quantity and quality of pollen acquired by honeybees positively influences their health and survival by improving their longevity and immune function [2, 3]. Therefore, understanding the factors that affect honeybee pollen intake and foraging patterns is fundamental to enhancing their well-being and productivity.

Honeybees select among different pollen species according to their abundance in the surroundings, their attractiveness and the ease of collection and handling [4, 5]. In spite of this selectivity, however, honeybees forage on diverse resources, including some that are not particularly attractive or efficiently gathered, in order to fulfill the great pollen demands required for successful colony development and reproduction (~20 kg year) [6], and to obtain different nutrients that may be segregated among plant species [2, 7]. Given such great pollen requirements, the amount and diversity of pollen collected by honeybees are expected to vary in response to changes in pollen availability due to agricultural intensification and flowering periods that are driven by climate seasonality. However, contradictory results have been found regarding the relationship between the amount and diversity of pollen and landscape structure in temperate agricultural regions [7, 8, 9, 10, 11], as honeybees seem to compensate for resource scarcity by increasing foraging distances to obtain sufficient quantities of diverse pollen for their colonies [10, 11, 12, 13]. However, temporal resource shortages resulting from flowering seasonality have also been found to considerably impact honeybee colony survival, especially in cases of landscapes with low diversity where pollen availability cannot be compensated for by extending foraging range [2, 10, 11].

How honeybees use pollen resources among different spatial and temporal conditions in tropical regions has been barely studied. Most such studies have been local and descriptive and, to the extent of our knowledge, there has been no study that explicitly related landscape structure and climate seasonality to the amount, richness and diversity of pollen collected by honeybees. This issue is of particular concern in the Neotropics, where foraging patterns of introduced honeybees have the potential to negatively impact native pollinators through competition and the pollination of native plant species [14]. Furthermore, significant losses of honeybee colonies are being reported in the Neotropics, as has been the case elsewhere in the world, while possible causes linked to floral resource scarcity prompted by agriculture intensification or global warming are still unknown [15].

The present work investigated the effects of landscape structure and climate seasonality on the amount, richness and diversity of pollen collected by honeybees in an agricultural region of Colombia where pollen production represents the principal income source for beekeepers, with yields of about 35 kg/hive/year [16]. Pollen species collected by honeybees in forest and anthropic areas (i.e., croplands, grasslands, woodlands and urban areas) were identified through palynological analysis. The amount, richness and diversity of pollen from forest species, anthropic species and overall were correlated with a gradient of landscape diversity and forest area, with the expectation of finding higher overall amounts of pollen with greater contribution of forest species in apiaries located in more diverse landscapes with larger forest areas, which are suspected to have greater resource availability. Temporal changes caused by climate seasonality were also explored by relating the amount and diversity of pollen from forest species, anthropic species and all species together with temporal variation in precipitation, given that rain is the most important climate driver for colony development and flowering phenology in tropical regions [17, 18]. The expectation was that precipitation would have negative effects since honeybees avoid flying during rain [19], and the flowering peaks of most species occur during dry weather conditions in this region [20]. Finally, the results were compared with those for temperate regions, and the possible implications for honeybee productivity and survival, and their interactions with native plants and pollinators, discussed.

## Materials and methods

### Study region

This work was performed in an agricultural region of Colombia (elevation 2500–3000 masl), in the surroundings of the city of Bogotá. The land cover was represented by a mix of cropland, grassland, woodland and forest fragments of various sizes (Low and High Andean Forest Ecosystems) [21], with the cultivation of potato, pine and eucalyptus accounting for most of the cropland and woodland area. The climate is characterized by a relatively stable temperature throughout the year (mean annual temperature of 9–14°C, monthly variation coefficient 2.3%) and a bimodal distribution of precipitation with rainy periods from March to May and from September to November (annual precipitation of 600–1500 mm, monthly variation coefficient 42.3%) [22].

### Landscape and climate data

Ten apiaries were selected according to a landscape gradient of forest cover, from 2% to 78% in buffers of 1000m around each apiary [13, 23] (site coordinates summarized in S1 Appendix). Areas of forest and anthropic cover (i.e., croplands, grasslands, woodlands and urban areas) were calculated by creating polygons on Landsat images (scale 1:10000) from Google [24] using QGis 2.18.0 (GNU-GPL, Boston, MA). Landscape diversity was estimated by the Shannon Diversity Index with the proportional areas of cover types [25]. Data of mean monthly precipitation were obtained from the database of Instituto de Hidrología, Meteorología y Estudios Ambientales for a hydrological station in Bogotá [22].

### Pollen data

Pollen samples (500 g) were collected from Africanized honeybee hives [26], once a month between September 2009 and March 2010, using modified Ontario Agricultural College pollen traps [27]. Samples were obtained from bulk containers where pollen from all hives in the apiary was mixed together after harvested by beekeepers. Data regarding pollen harvest weight and number of harvested hives were recorded for each sampling journey.

Pollen samples were prepared through acetolysis [28] and mounted on slides for grain counting and species identification under a microscope. The slides were deposited in the palynological collection of the Laboratorio de Investigaciones en Abejas, Universidad Nacional de Colombia. Pollen amount was calculated for each pollen morphotype by multiplying grain relative abundances by mean grain volume for the morphotype [29]. Proportional amounts were then calculated and used to classify pollen types as: dominant (> 45%), secondary (>15–45%), important–isolated (3–15%) and isolated (< 3%) [30].

Pollen identification was performed by P.M.M.P., to the lowest taxonomic level possible, with the help of reference slides obtained from plants collected in the field and pollen catalogs [31, 32, 33]. Identified morphotypes were classified according to their habitat as either forest species (those species originally from forest areas but could also be found in both forest and anthropic areas) or anthropic species (those species strictly associated with anthropic areas). Unidentified pollen morphotypes were dropped from the analysis since information could not be obtained about their habitat (18 pollen types, 1.36% of total pollen amount).

### Statistical analysis

Data were analyzed using generalized linear models (GLM). The amount, richness and diversity of all pollen species and forest species and anthropic species separately, where taken as response variables, whereas the main effects and the interaction between forest area, landscape diversity and monthly mean precipitation were used in the initial predictor model. The best models were selected based on significant differences in AIC values (ANOVA tests, significance level = 0.05), while residuals were visually inspected to assess model fit. All analyses and figures were performed in R v.3.5.0 software [34], using the packages MASS [35], vegan [36], bipartite [37], ggplot2 [38], gridExtra [39] and ggsci [40].

## Results

A total of 103 pollen morphotypes distributed in 22 families were found in the pollen samples, of which 47 were classified as forest species (9% of overall pollen amount) and 41 as anthropic species (89% of overall pollen amount), while 18 could not be classified as they could not be identified to the species level (1.36% of overall pollen amount). Nine species were found in more than 50% of the samples, and comprised 72% of all pollen. These species included the anthropic species *Brassicaceae* type namely *Brassica rapa* and *Raphanus raphanistrum* with undifferentiated pollen morphology (97% of samples, mean pollen amount 23 ± 16%), *Hypochaerys radicata* (97% of samples, mean pollen amount 14 ± 12%)*, Eucalyptus globulus* (87% of samples, mean pollen amount 8 ± 8%), *Trifolium pretense* (81% of samples, mean pollen amount 14 ± 13%) and *Trifolium repens* (78% of samples, mean pollen amount 6 ± 6%), as well as the forest species *Rubus floribundus* (60% of samples, mean pollen amount 4 ± 5%), *Abatia parviflora* (59% of samples, mean pollen amount 1 ± 1%)*, Muehlenbeckia tamnifolia* (54% of samples, mean pollen amount 1 ± 1%) and *Baccharis* (51% of samples, mean pollen amount 1 ± 1%) (Fig 1). Families with the greatest amount of species were Compositae with 14 species, Leguminosae with four species, and Adoxaceae, Myrtaceae, Polygonaceae, Rosaceae and Solanaceae with two species each.

**Fig 1.**
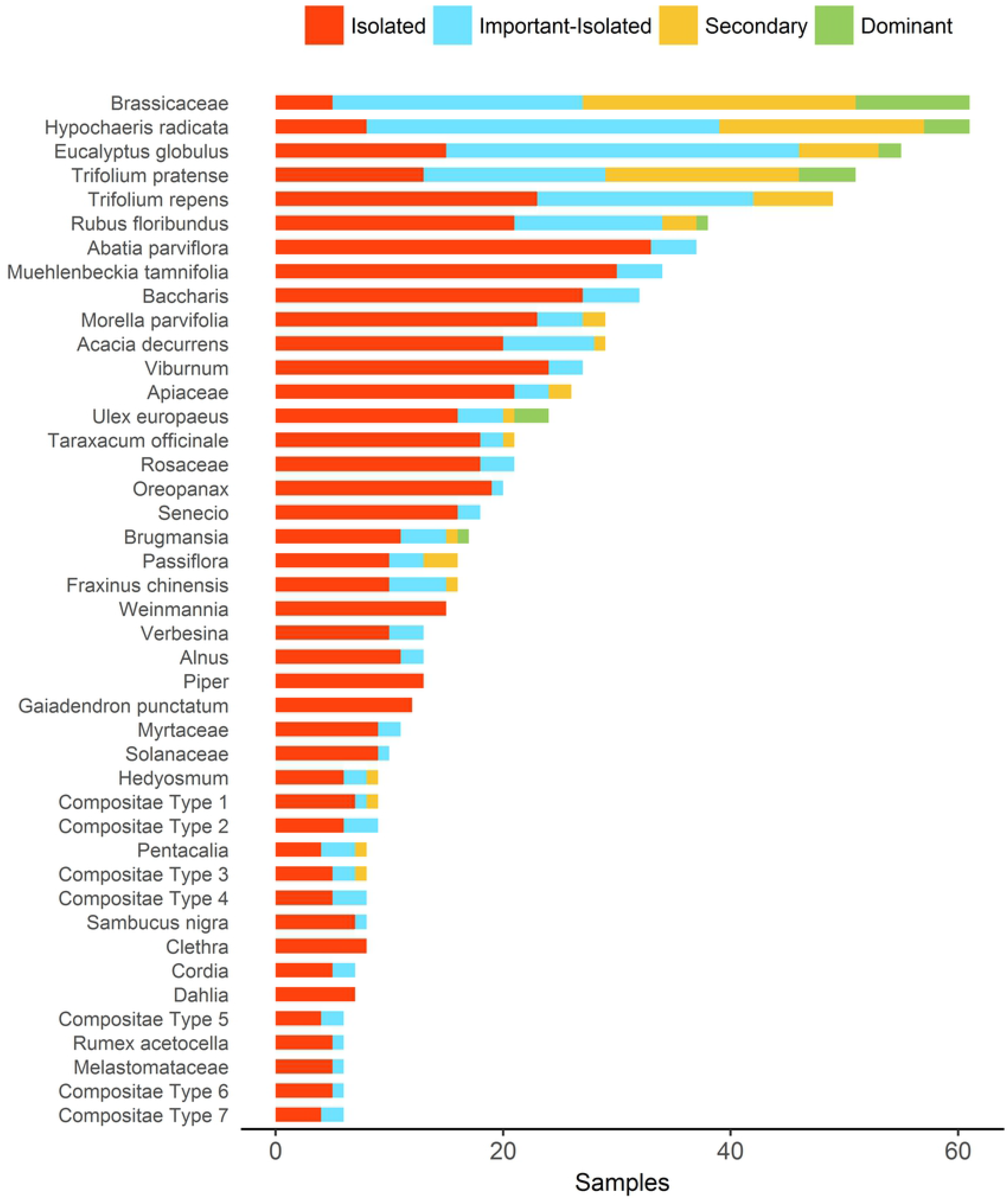
Most important pollen species foraged by honeybees in an agricultural region in Colombia. The figure shows 43 out of 103 species found in pollen samples, which comprise 95% of total pollen amount and were found in more than 10% of pollen samples.

The overall pollen amount collected by honeybees mainly corresponded to anthropic species (Fig 2a), and was positively related to forest area (explained variance = 0.04, p-value < 0.001, Fig 3a, Table 1), landscape diversity (explained variance =0.04, p-value < 0.001, Fig 3b) and the interaction between these two variables (explained variance = 0.24, p-value < 0.001, Fig 4a). The pollen amount from forest species was positively affected by forest area (explained variance = 0.07, p-value = 0.03, Fig 3a) and landscape diversity (explained variance = 0.05, p-value = 0.07, Fig 3b), while the pollen amount from anthropic species was negatively affected by forest area (explained variance = 0.07, p-value=0.03, Fig 3a) and landscape diversity (explained variance = 0.05, p-value=0.07, Fig 3b).

**Fig 2.**
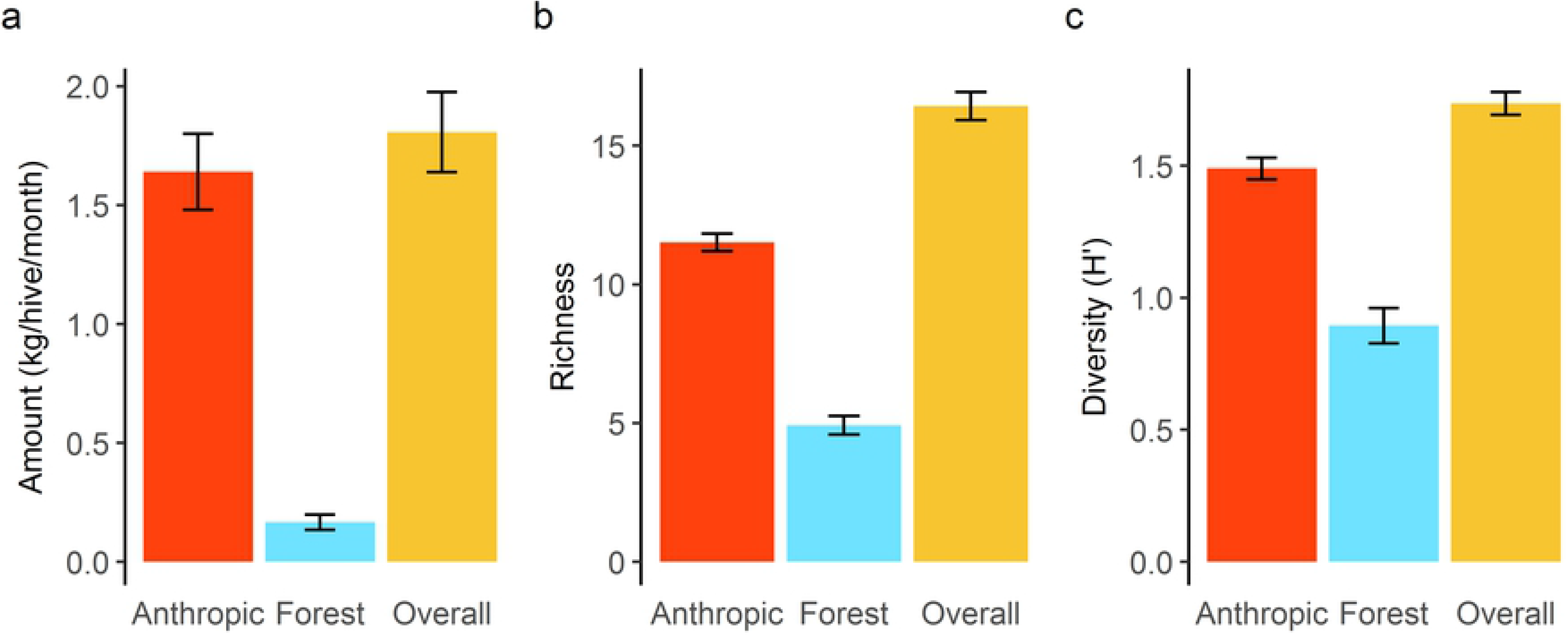
Mean amount (a), richness (b) and diversity (c) of pollen collected by honeybees. Colored bars indicate pollen from species found in the overall landscape around the apiary (1000 m radius buffers) and segregated into pollen from anthropic areas (i.e., croplands, grasslands, woodlands and urban areas) and from forest areas. Error bars indicate standard error values.

**Fig 3.**
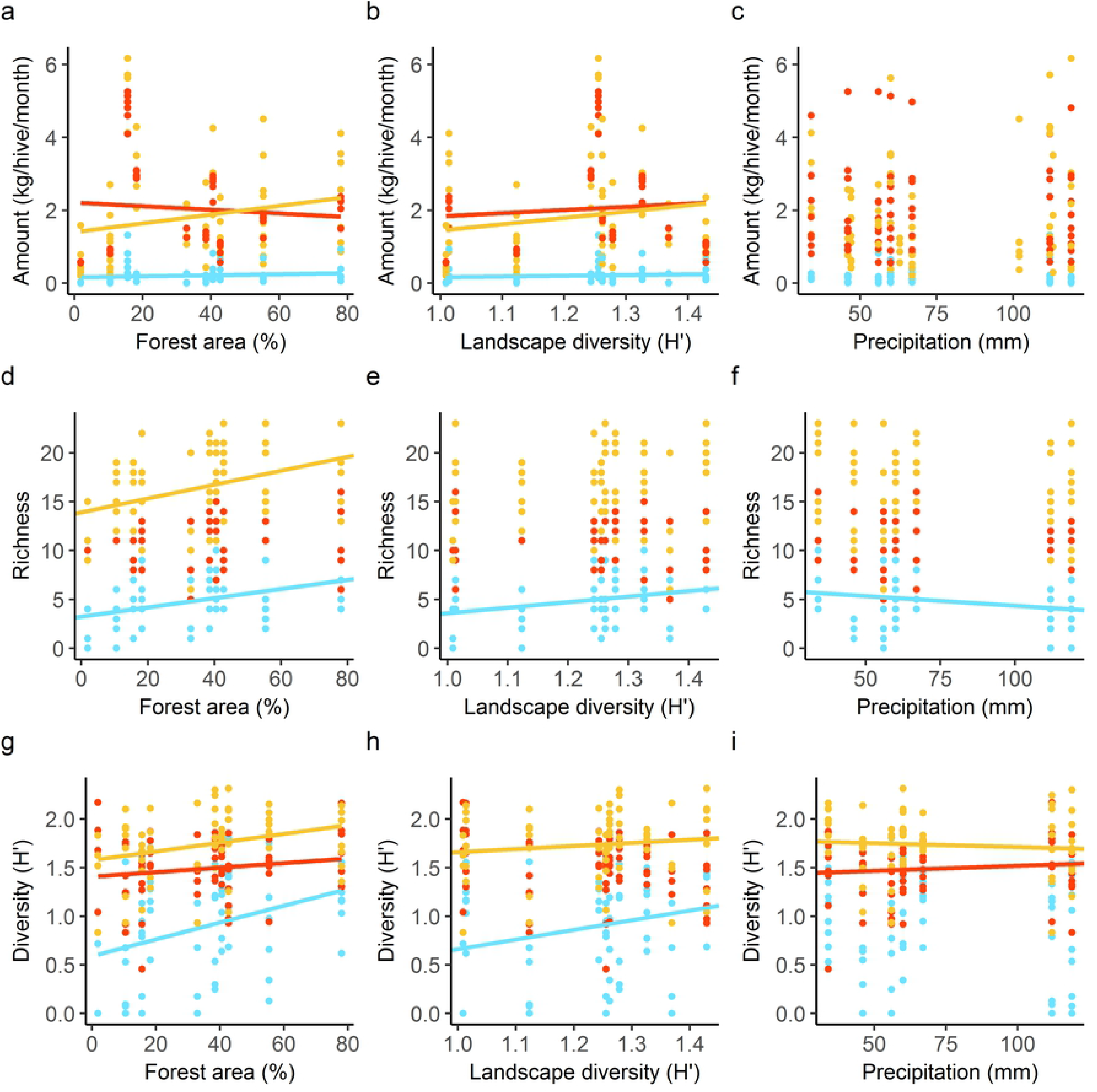
Main effects on the amount, richness and diversity of pollen collected by honeybees. Forest cover area (a, d, g), landscape diversity (b, e, h) and mean monthly precipitation (c, f, i) were taken as main effects in generalized linear models to explain the amount, richness and diversity of pollen collected by honeybees from species in the overall landscape (yellow dots), and from species segregated as from forest areas (blue dots) and anthropic areas (red dots). Significant effects are represented by tendency lines in the color of the respective response variable (confidence level 0.05).

**Table 1.** Selected models explaining variation in the amount, richness and diversity of pollen collected by honeybees.

| Response variable | Selected model | R<sup>2</sup> / D* | p-value |
| --- | --- | --- | --- |
| Overall amount | $- 5.69 + 0.17FA + 9.26LD - 0.21FA * LD$ | 0.29 | <0.001 |
| Forest amount | $- 116.66 + 0.12FA + 17.35LD$ | 0.09 | 0.019 |
| Anthropic amount | $- 116.66 - 0.12FA - 17.35LD$ | 0.09 | 0.019 |
| Overall richness | $14.07 + 1.00FA$ | 0.14 | NA |
| Forest richness | $- 0.81 + 1.01FA + 3.72LD - 1.00P$ | 0.25 | NA |

| Anthropic richness | NULL |  |  |
| --- | --- | --- | --- |
| Overall diversity | $3.63 + 0.004FA - 1.62LD - 0.03P + 0.03LD * P$ | 0.14 | 0.011 |
| Forest diversity | $- 0.67 + 0.008FA + 1.02LD$ | 0.17 | 0.002 |
| Anthropic diversity | $0.96 + 0.01FA + 0.006P - 0.0001FA * P$ | 0.07 | 0.056 |

Response variables were the amount, richness and diversity of pollen from species in the overall landscape, and from species segregated as from forest areas and from anthropic areas. Main effects of the initial models included the interaction between forest area (FA), landscape diversity (LD) and mean monthly precipitation (P). Best models were selected by ANOVA testing of AIC values. \*Explained deviance (D).

**Fig 4.**
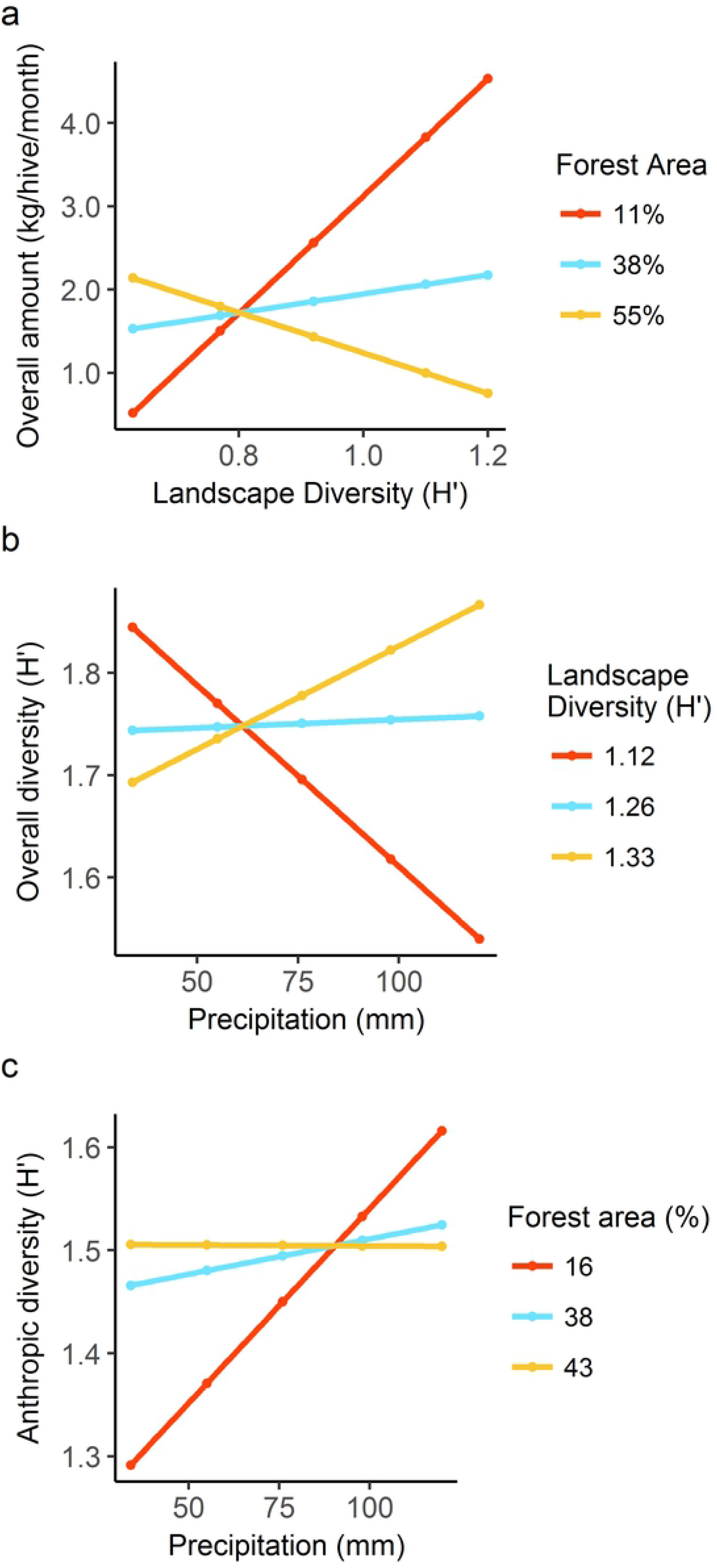
Main-effect interactions explaining the amount and diversity of pollen collected by honeybees. (a) Interaction between landscape diversity (Shannon’s H’) and forest cover area explaining variation in overall pollen amount. (b) Interaction between mean monthly precipitation and landscape diversity explaining variation in overall pollen diversity. (c) Interaction between mean monthly precipitation and forest area explaining variation in pollen diversity from species in anthropic areas. Legend numbers correspond to 25%, 50% and 75% quantiles in the distribution of the respective variable.

Overall pollen richness was dominated by anthropic species (Fig 2b) and was explained by forest area (explained deviance = 0.14, p = 0.002, Fig 3d). Forest pollen richness was positively related to forest area (explained deviance = 0.13, p = < 0.001, Fig 3d) and landscape diversity (explained deviance = 0.09, p = 0.003, Fig 3e), while negatively affected by precipitation (explained deviance 0.03, p = 0.066, Fig 3f). Anthropic pollen richness was not explained by any of the studied variables.

Overall pollen diversity mainly corresponded to anthropic species (Fig 2b), and was explained by forest area (explained variance = 0.08, p=0.03, Fig 3g), landscape diversity (explained variance = 0.02, p = 0.05, Fig 3h), precipitation (explained variance = 0.001, p = 0.01, Fig 3i), and the interaction between precipitation and landscape diversity (explained variance = 0.09, p = 0.01, Fig 4c). Forest pollen diversity was positively affected by forest area (explained variance = 0.13, p = 0.002, Fig 3g) and landscape diversity (explained variance = 0.06, p = 0.03, Fig 3h). Anthropic pollen diversity was positively affected by forest area (explained variance = 0.02, p = 0.01, Fig 3g), precipitation (explained variance = −0.01, p = 0.02, Fig 3i) and the interaction between these two variables (explained variance = −0.08, p = 0.02, Fig 4c).

## Discussion

This work analyzed the effects of landscape structure on the amount, richness and diversity of pollen collected by honeybees, with the expectation that there would be greater overall pollen amounts with higher contributions of forest species in more diverse landscapes with larger forest areas. Contrary to this expectation, honeybees were found to be principally associated with anthropic species despite any variation in landscape structure (Fig 2). Such a high preference might be because anthropic plants mainly correspond to weeds and crops, which flower massively in dense patches. These species, therefore, could be more easily foraged by honeybees than the sparse populations of forest species [41], which in turn may require higher efforts to fly further between conspecific plants and to waggle-dance to communicate resource locations [42, 43, 44]. Nonetheless, several cases of species with dense populations in more conserved tropical forests (e.g., *Quercus humboldtii* and *Weinmania tomentosa* in mature Andean forests and *Rizophora mangle* in mangroves), have also been registered as predominant species in honeybee pollen samples [45, 46].

The most important anthropic species (i.e., Brassicaceae Type, *Trifolium* species and *H. radicata* Fig 1) also corresponded to invasive alien plants that have interacted and coevolved with honeybees for a much longer period of time than Neotropical native plants [47]. Besides, alien species flower more constantly than forest species since the environmental constraints that mediate their flowering periods in temperate areas, such as seasonality in temperature and day length [48], are limited in this tropical region. Additional constraints among forest species could also have hindered pollen foraging by honeybees, such as poricidal anthers in the families Melastomataceae and Solanaceae, for example — families which also possess many species that are dominant in tropical forests [21]—, which require vibrational movements to extract pollen; an activity that cannot be executed by honeybees [49].

In spite of the greater importance of anthropic species, the present results demonstrated that the amount, richness and diversity of pollen from forest species and from all species together were positively related to the proportion of forest area around the apiaries, as originally predicted (Fig 3a, d, g). These results suggested an advantage for honeybees in obtaining small quantities of pollen from diverse resources in forest areas, which is probably related to a gain in nutritional content and the subsequent health benefits [2, 7].

Landscape diversity was also positively related to the amount, richness and diversity of pollen from forest species and overall species, probably as a result of larger forest areas associated with the complex landscape. However, landscape diversity was also positively related to the amount of pollen from anthropic species, suggesting that the honeybees also benefited by an increase in diversity of anthropic cover types. The interaction between landscape diversity and forest area revealed that the proportion of forest area became detrimental to the overall pollen amount when it surpassed a threshold of about 40% of total land cover (Fig 4a), seemingly corroborating the idea that honeybees rely principally on anthropic species even when forest species could represent an additional advantage.

Similar correlations between landscape/resource diversity and pollen diversity collected by honeybees were found in a subtropical region of the United States; [8] and in western France [7]. However, various studies in temperate regions of Europe [9, 11] and the United States [10] found no relationship between the amount and diversity of pollen and landscape diversity, since honeybees compensated for less diverse landscapes with small proportions of natural and semi-natural areas by increasing their foraging range and focusing scouting activities on few pollen resources, thus maintaining the amount and diversity of the collected pollen [11, 12, 13, 50]. Such compensation was probably not required in the present study because less diverse landscapes with small forest areas still provided several anthropic species to sustain honeybee colonies throughout the year, while more complex landscapes appeared to stimulate honeybees to explore new resources, with a resultant gain in pollen amount.

Regarding climate variability, no relationship was found between precipitation and the amount of pollen collected by honeybees (Fig 3c), probably because monthly differences in precipitation were not important in determining their pollen foraging patterns. Studies carried out in tropical regions of the Americas [51] and Africa [17] have also found that honey bees forage on pollen and exhibit brood-rearing activity throughout the year in spite of climate seasonality, and that honeybees increase their foraging efforts the day before heavy rainfall [52], thus compensating for low activity during rainy days. Nonetheless, a negative effect was found for precipitation on the richness of pollen from forest species and the diversity of pollen from all species (Fig 3f, i), which may be explained by factors such as decreased flight activity [19], washing of pollen from flowers, and flowering shortage during rainy periods [20]. The interaction between precipitation and landscape diversity demonstrated, though, that overall pollen diversity decreased only in landscapes with low diversity, while it tended to increase in complex landscapes (Fig 4b). Studies in temperate regions have revealed that pollen diversity increases during periods of resource scarcity (e.g., late summer and fall), since honeybees switch to forage on a greater diversity of species to fulfil their pollen requirements, as individual species are less abundant and hence more quickly depleted [10, 11, 13; 50]. However, such compensatory behavior is not possible in extremely simple landscapes given that their lower resource availability cannot support more diverse pollen diets [11]. Similar to temperate regions, it is possible that the complex landscapes of the present work buffered the negative impacts of precipitation through higher resource availability, while the low diversity landscapes reinforced the negative effects of precipitation by increasing resource scarcity during rainy periods. Still, precipitation effects should not have excessively compromised honeybee survival since the amount of pollen collected by honeybees was not affected by precipitation.

As with the effect on overall pollen diversity, pollen diversity from anthropic species was found to be positively related to precipitation (Fig 3i), although the interaction between precipitation and forest area showed that the increase in anthropic pollen diversity occurred just in landscapes with small forest areas (Fig 4c), where the availability of anthropic cover types was probably sufficient to provide more diverse anthropic species. In contrast, anthropic pollen diversity in forested landscapes tended to decrease, probably because smaller anthropic areas hindered pollen gathering on diverse anthropic resources.

In conclusion, we found that honeybees in this Neotropical region were especially associated with plant species that are abundant in anthropic areas, and minimally associated with native forest species. These results suggest that competition between honeybees and native bees for floral resources would be minimal since their niche overlap was probably narrow, and would not cause considerable effects on pollination rates and gene flow of forest species as they were little visited. However, these assumptions should not be generalized given the opposite results found in other Neotropical regions [14], and thus deserve further research.

Landscape diversity and forest area were found to benefit honeybees by increasing pollen amount, richness and diversity. Therefore, beekeepers should consider locating their hives in mixed landscapes containing forest patches and diverse anthropic areas in order to enhance pollen production for honeybees. Intensification of agriculture did not seem to be an important cause of resource depletion for honeybees. However, other factors related to agriculture, such as use of pesticides, for example, should be taken into account and evaluated as possible key drivers of honeybee mortality in tropical agricultural regions [15].

Precipitation did not affect the amount of pollen collected by honeybees, although it affected pollen diversity, with stronger negative effects in landscapes with less resource availability. The predicted increase in precipitation with global warming in this region [53] could reinforce negative effects on pollen diversity and ultimately pollen amount by increasing the number of days without flight activity and the death of foragers caught in heavy rain, or indirectly by altering flowering phenologies or daily periods of anthesis [19, 54]. Future studies should contemplate longer sampling periods to test yearly differences in precipitation regimes, as well as waggle-dance decoding to go into honeybee responses to environmental changes in greater depth and to be able to make better comparisons with works in temperate regions like those of [11, 12, 13].

## Acknowledgments

We thank the beekeepers Yaneth Rondón, David Guzmán, Maria Fernanda Jimenez Alejandra Faraco and Alfonso Franky for providing us with pollen samples; the staff of Laboratorio de Investigaciones en Abejas LABUN for their support with palynological processing and identification; and Rodulfo Ospina Torres, Carlos Sarmiento and Marisol Amaya Marquez for their valuable suggestions.

## Supplementary material

**S1 Appendix. Original data of pollen samples obtained from honeybee hives.** Includes information about location, sampling date, pollen species composition, pollen richness and diversity calculations, proportion of forest area around apiaries (1000m buffers), landscape diversity (Shannon Index), and mean monthly precipitation (mm).

**S2 Appendix. Pollen amounts of forest, anthropic and all species collected by honeybees** Pollen quantities are expressed as per hive per month. Data for the explanatory variables of forest area, landscape diversity and mean monthly precipitation are included for running linear regressions.

